# Predicting semantic segmentation quality in laryngeal endoscopy images

**DOI:** 10.1101/2024.11.14.623604

**Authors:** Andreas M. Kist, Sina Razi, René Groh, Florian Gritsch, Anne Schützenberger

## Abstract

Endoscopy is a major tool for assessing the physiology of inner organs. Contemporary artificial intelligence methods are used to fully automatically label medical important classes on a pixel-by-pixel level. This so-called semantic segmentation is for example used to detect cancer tissue or to assess laryngeal physiology. However, due to the diversity of patients presenting, it is necessary to judge the segmentation quality. In this study, we present a fully automatic system to evaluate the segmentation performance in laryngeal endoscopy images. We showcase on glottal area segmentation that the predicted segmentation quality represented by the intersection over union metric is on par with human raters. Using a traffic light system, we are able to identify problematic segmentation frames to allow human-in-the-loop improvements, important for the clinical adaptation of automatic analysis procedures.

## Introduction

Semantic segmentation is the pixel-wise classification of present objects in a given image. Especially in a medical context, semantic segmentation of different tissues is an important task and has seen major improvement since the advent of Deep Learning [1]. In contrast to object detection, semantic segmentation delineates specific organs and allows the precise quantification of covered areas. In the case of laryngeal endoscopy, the area between the vocal folds, the so-called glottal area, is an important proxy for the vocal folds’ oscillation behavior [2, 3]. Many works have used a variety of classical and Deep Learning-based computer vision techniques to approach glottal area segmentation [4], ideally in a fully automatic manner [5–7]. A plethora of clinical parameters have been described [8] that rely on the glottal area waveform, a biosignal derived from the glottal area. As the validity of these parameters heavily depends on the segmentation quality of the glottal area [9], it is important to assess this quality using meaningful metrics.

Common metrics to evaluate this segmentation quality are the Dice and the Intersection over Union (IoU) score. Both scores compare the ground-truth area and predicted area and compute a score between 0 and 1 that needs to be maximized. Interestingly, Dice and IoU scores are highly related and can be converted into each other by a simple math formula [10], hence, we use for consistency only the IoU score in the following. However, both need the underlying ground-truth segmentation commonly provided by domain experts. Providing this often manually generated ground-truth data in a clinical context is not feasible, as it involves a significant amount of hands-on time [7]. Therefore, previous studies proposed an AI-powered system to predict the IoU/Dice score of a given segmentation to hint at potential failure cases [11, 12]. Such a system for analyzing high-speed videoendoscopy footage segmentation that typically consists of 4,000 frames per second acquisition time would be highly beneficial but is nonexistent.

However, the IoU score itself has no contextual meaning about its clinical usefulness raising the issue which IoU scores are actually desired. We agree that an IoU score of 1 is thought to be perfect, but which IoU scores can be reached unequivocally by domain experts? First evidence has been given in [7] showing that the average IoU score for glottal area segmentation for three experts is 0.772, a value reasonably distant to 1.

However, Maryn and colleagues [13] have demonstrated that the glottal area segmentation across individuals on that quality level barely affects the downstream parameter computation. However, inter- and intra-rater reliability has been not assessed clearly for glottal area segmentation.

In this study, we address these issues by providing an in-depth analysis of inter- and intra-rater reliability and developing a novel IoU prediction system for glottal area segmentation. With this information, we suggest a traffic light system for glottal area segmentations to pinpoint problematic frames observed during the analysis to guide the clinician in interpreting downstream computed parameters with respective care.

## Materials and methods

### Data

In this study, we utilized the Benchmark for Automatic Glottis Segmentation (BAGLS, [7]) for analyses presented in Fig. 1-3. The data used in Fig. 4 consisted of retrospective medical records accessed on February 15, 2024. The individual authors did not have access to the subject identities during data access. We used custom Python scripts to create the target dataset (see Github). We cropped images of an aspect ratio different from 1:1 around the glottal area and resized each frame to a common resolution of 224 *×* 224 *×* 3. This training data is available through Zenodo (see Data Availability). The ground-truth masks were modified to generate segmentation masks with segmentation artifacts. The full process is shown in Figure 1. In particular, we used erosion and dilation to modify the mask uniformly. For uncertainties at the glottal area border, a Sobel filter was applied to detect the edges. After binary dilation, each edge pixel was randomly set to be added or removed from the ground truth. Small segmentation artifacts close to the glottis were mimicked using up to five randomly drawn spheres with a random radius between one and three pixels. For large segmentation artifacts, 2D Perlin noise was added to the ground truth mask. Here, we relied on the Perlin noise generator for numpy^1^.

**Fig 1.**
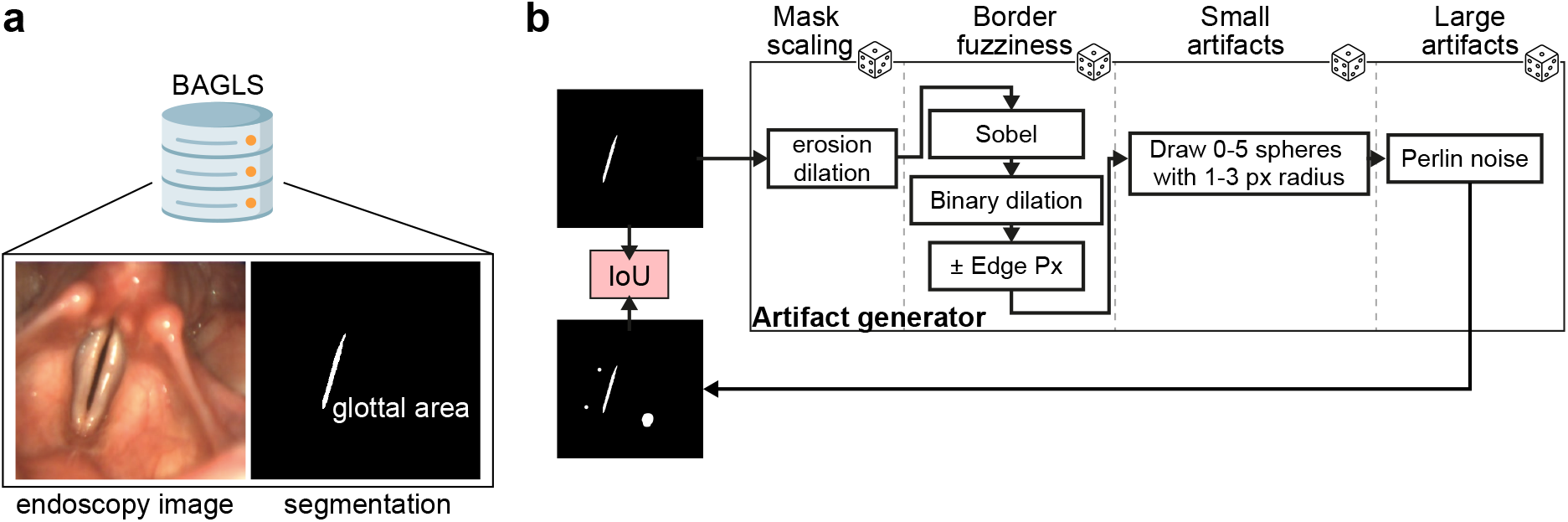
Artifact generation process. A: The BAGLS dataset contains 55,750 paired training samples consisting of endoscopy images (left) and their respective glottal area segmentation (right). B: Ground-truth glottal area segmentation is sent to the artifact generator. We apply four steps to incorporate uniform mask scaling artifacts, border fuzziness, and small and large segmentation artifacts. Each step is randomly applied, and step-dependent hyperparameters are randomly chosen. The resulting segmentation mask is used to compute the IoU score with the ground-truth segmentation. The resulting segmentation masks together with the IoU score are used for training downstream deep neural networks.

**Fig 2.**
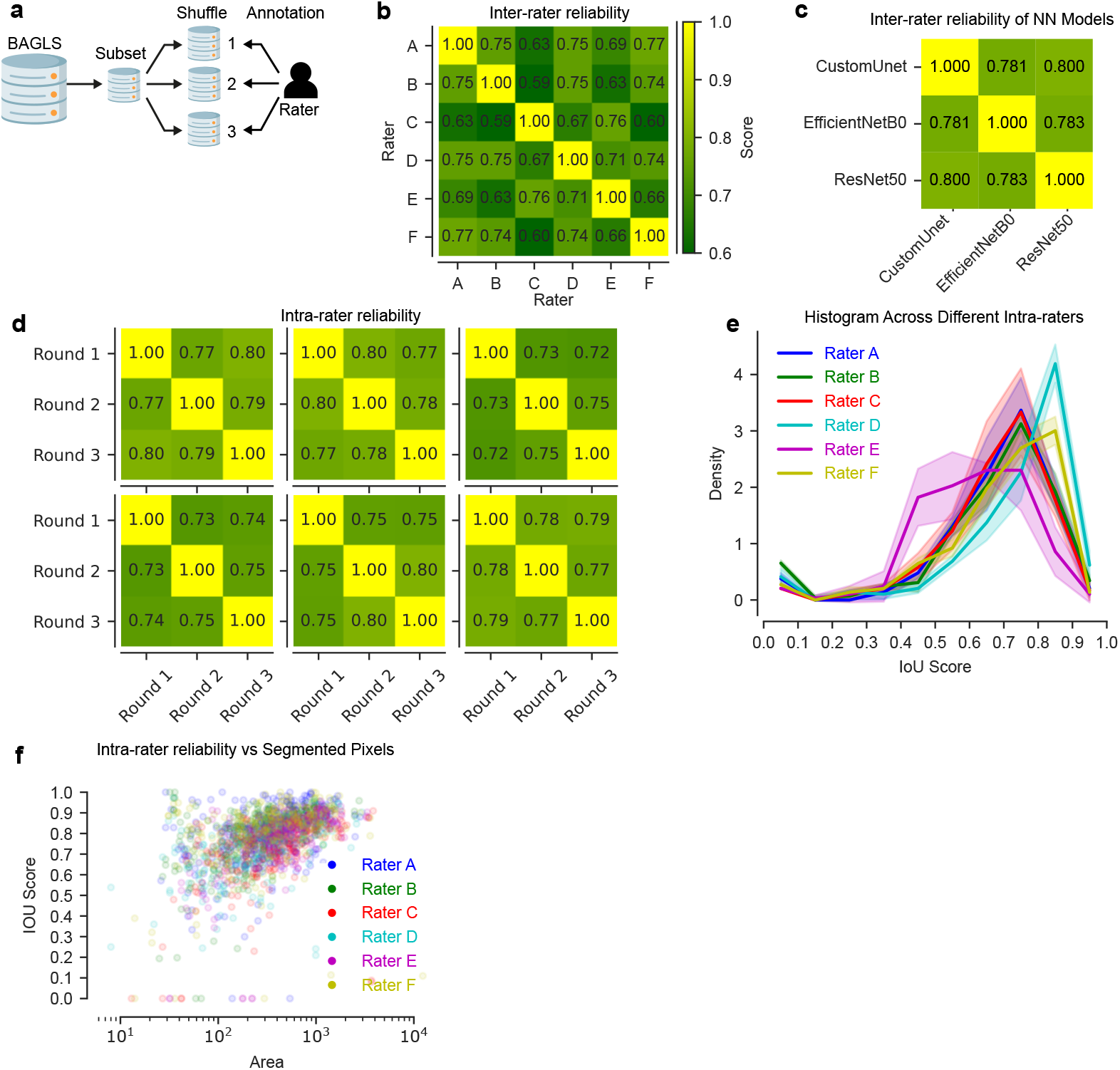
Inter- and intra-rater reliability shows non-perfect agreement. (A) Task overview. A subset of the BAGLS dataset was taken (100 random frames) and annotated by trained raters three times in random order. (B) Details inter-rater reliability among six raters, highlighting consistency in their evaluations. (C) Presents the inter-rater reliability among neural network models described in [7]. (D) Explores intra-rater reliability for six raters across three rounds, assessing individual consistency. (E) Histogram across different Intra-raters. (F) Analyzes the relationship between inter-rater reliability and the number of pixels in segmented areas.

**Fig 3.**
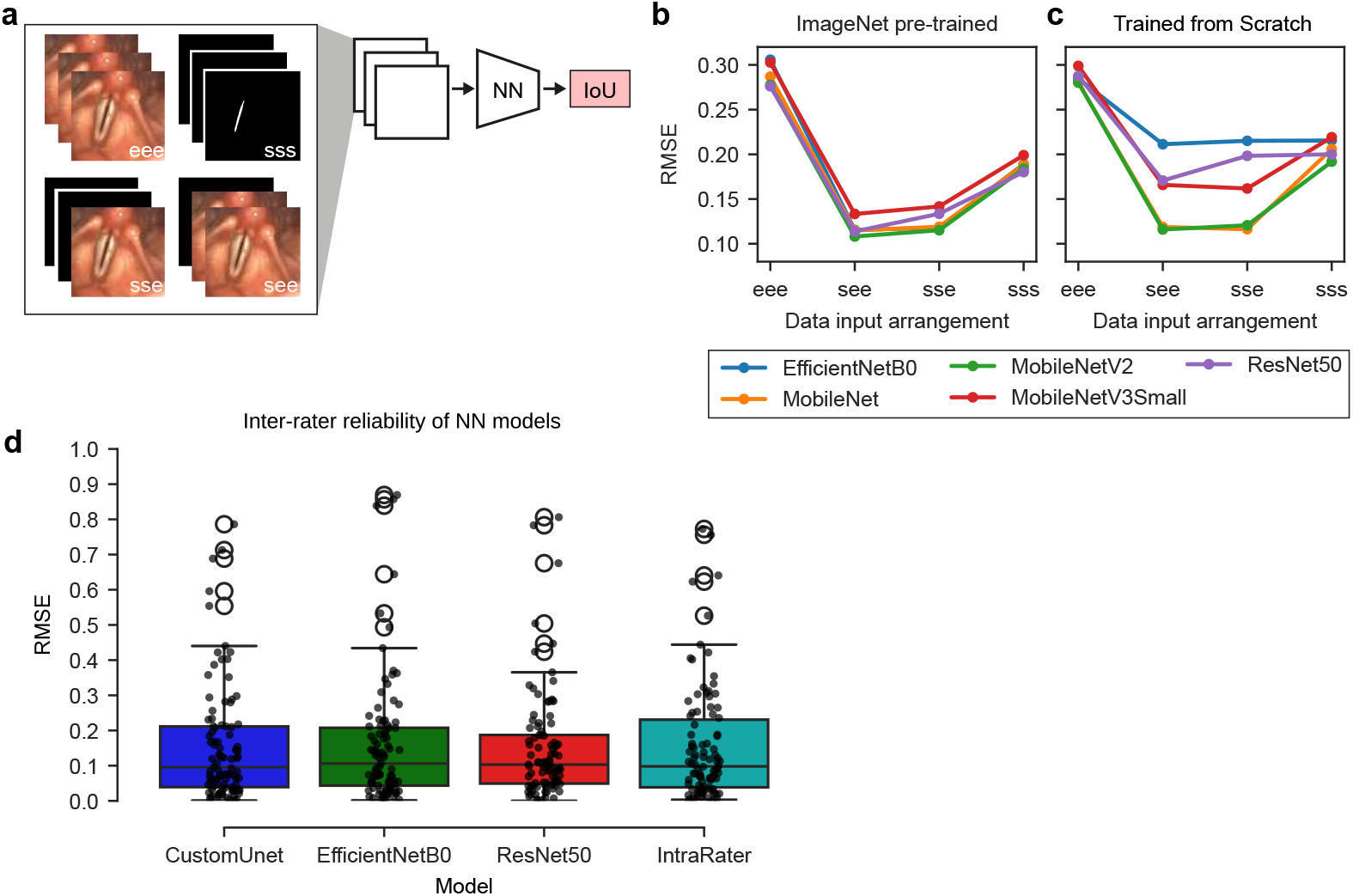
Segmentation quality prediction using neural networks. (A) Showcases various combinations of endoscopic images and their corresponding segmentation processed through the neural network architecture. (B, C) Illustrate outcomes across diverse network backbones under two distinct scenarios. (D) Comparison of RMSE between different neural networks and the average of six human raters.

**Fig 4.**
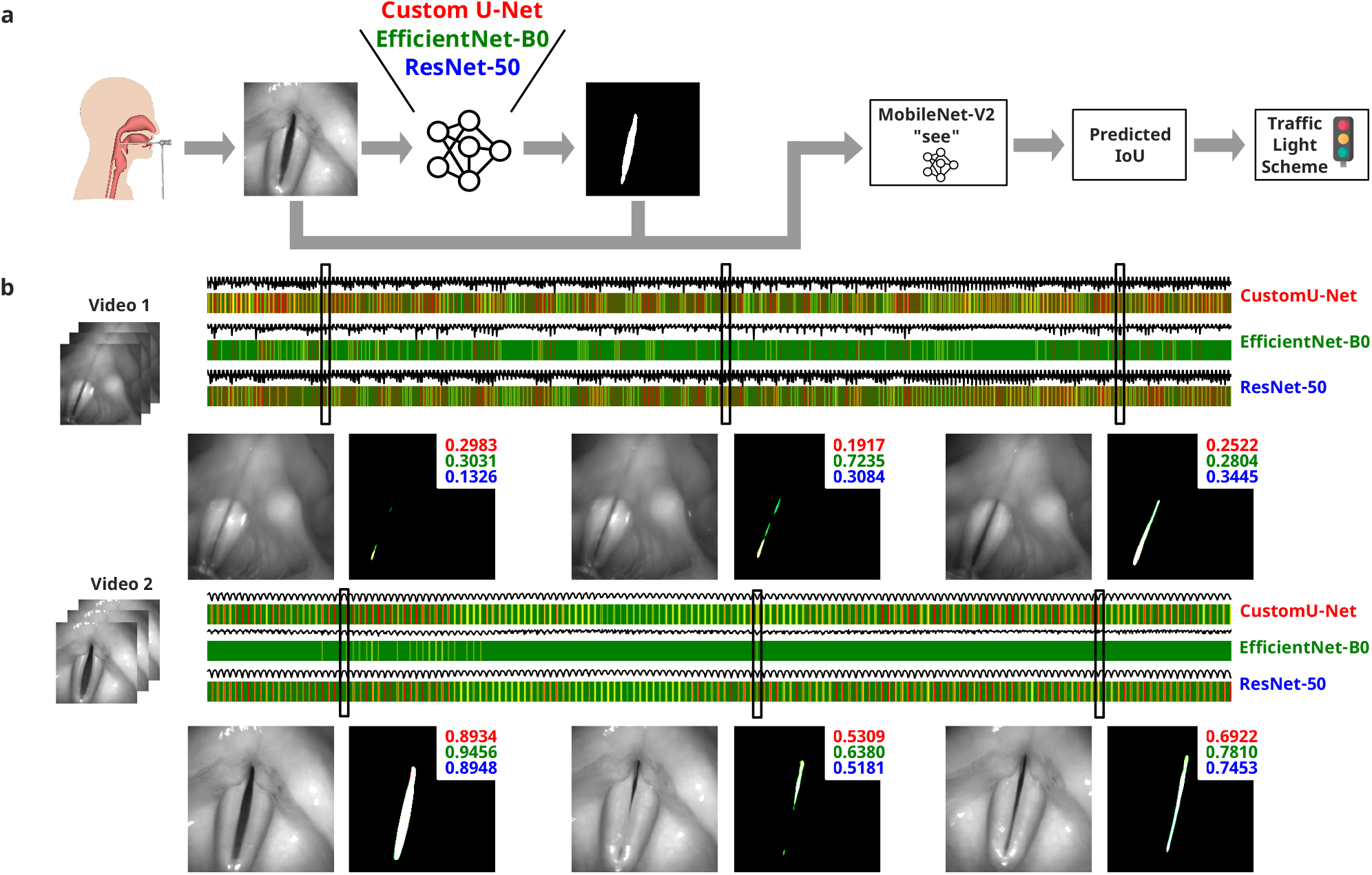
Traffic Light System. (A) Inference on independent test videos: For each frame, the glottal area is segmented using three different segmentation networks. The predicted masks and the original input frame are used as input (format “see”, as demonstrated in Fig. 3b/c) to the trained MobileNetV2, which predicts the IoU score. A traffic light scheme is applied to these results. (B) Examples of traffic light bars using two exemplary videos: The traffic light bars show the predicted color for each segmentation network. For each video, three exemplary frames and their corresponding IoU predictions are shown.

The Intersection over Union (IoU) score is our target metric (see evaluation) and ranges from 0 (no overlap, bad segmentation) to 1 (perfect overlap, excellent segmentation). We used 20 discrete bins each with a width of 0.05 to ensure an even sampling across the full IoU range. From the BAGLS dataset, we drew random frames to generate 2,000 training pairs in each bin to ensure a balanced dataset. For each image, we applied randomly combinations of the above-mentioned techniques to the segmentation mask and determined the IoU score compared to the untreated ground-truth segmentation mask.

### Deep Neural Networks

All deep neural networks were set up using TensorFlow/Keras in version 3.1.1 of Keras and 2.16.0 of Tensorflow with enabled CUDA/cuDNN support. Experiments were performed on an RTX A4000 GPU. We relied on established architectures that have been widely used across tasks. Specifically, we utilized the MobileNet, MobileNetV2, MobileNetV3, ResNet50, and EfficientNetB0 [14–18] architecture to regress the IoU score. In detail, we used the backbone architecture with their respective `tensorflow.keras.applications` implementation removed the top classification layer, added a GlobalAveraging2D pooling layer, then a Dense Layer with 256 units and ReLU activation, and a Dense layer with a single unit and sigmoid activation function. For regularization, a Dropout Layer with 0.1 dropout probability was placed between the two Dense layers. Neural networks were trained for 50 epochs using either SGD with Momentum (set to 0.9) or the Adam optimizer and an initial learning rate *η* of 10^−2^ and 10^−3^, respectively, with a reduction of *η · e*^−0.1^ after 10 epochs. We evaluated the mean squared error (MSE), the mean absolute error (MAE) and the ordinary categorical cross-entropy as a loss function, where the latter was used for subsequent analyses. We either provided the endoscopy frame (*e*) or the segmentation mask (*s*) as input in various combinations to predict the target IoU score. The combinations are (i) only endoscopy images (abbreviated as eee), (ii) only segmentation masks (sss), (iii) two endoscopy images and one segmentation mask (ees or see), (iv) two segmentation masks and one endoscopy image (sse or ess). We used three channels for convenience in training, as pre-trained networks expect RGB images with three channels. For our experiments, any RGB endoscopy image was converted to grayscale prior to training. Input frames were normalized to a range from -1 to 1.

### Inter- and intrarater analysis

The critical importance of assessing inter-annotator agreement is emphasized for fundamental reasons [19]. With high inter-rater reliability (IRR), the consistency of annotations, crucial for designing robust AI algorithms, is ensured together with effective AI model training and evaluation by reducing noise and subjectivity. It further contributes to overall annotation validity, signifying good data quality for reproducible studies.

Therefore, in our paper, six independent annotators manually annotated the glottal area in the same 100 randomly selected frames from the BAGLS dataset exactly three times. Each annotation round consisted of the exact 100 frames but randomly shuffled in order to avoid any progression bias. For glottis annotation, we used the Pixel Precise Annotator (PiPrA, [7]) to generate single segmentation masks for each frame individually.

### Prediction of segmentation quality using Traffic Light System

Five videos of healthy subjects containing at least 2,000 frames were analyzed. Each frame was segmented using previously established glottis segmentation networks [8], including CustomUnet *Fast* [7], ResNet50 *Quality* and EfficientNetB0 *Balanced* [20], and the segmentation quality, i.e. the IoU score, was predicted. For IoU score prediction, we relied on the MobileNetV2 S-E-E variant shown in Figure 3. The reason for selecting this variant is thoroughly explained in the Results section.

Each video is visualized by a bar whose colors are derived from a traffic light scheme computed for each of the 2,000 frames. Each frame’s IoU score follows the following color code: green indicates high IoU scores (above 0.7), suggesting good segmentation quality as derived from the IeRR analysis; yellow signals moderate IoU scores (between 0.6 and 0.7); and red denotes low predicted IoU scores (below 0.6), highlighting areas of potential concern. These values are in line with our IeRR study and with recent reports [21]. This visual tool allows for an immediate evaluation of the performance of various deep neural network models over continuous video frames.

### Evaluation

The common metric to evaluate the semantic segmentation quality is the Intersection over Union (IoU) and the Dice score. These metrics have been assessed in many works previously [6–9, 21]. As IoU and Dice scores are mathematically related (e.g. [22]), we solely report the IoU score in this work.

Inter-rater and intra-rater reliability (IeRR and IaRR) were assessed by comparing each rater to each other and its own three-fold annotations, respectively, using the IoU score. We determined the IaRR and IeRR by comparing each mask to each other (IaRR) or the average IoU score across masks (IeRR). Any error is shown as the standard error of the mean.

### Data availability

The code used to generate the annotation data, to generate the IoU prediction data, and to train the deep neural networks is available at https://github.com/ankilab/predicting_iou. The annotated frames for the intra- and inter-rater reliability study and the training data for IoU prediction can be found on https://doi.org/10.5281/zenodo.14034494.

## Results

### Intra- and inter-rater reliability

To find an automatic solution for segmentation quality prediction, we first need to investigate the variability and reliability across manual annotations by humans, the current gold standard in glottis segmentation. In total, six trained raters with various experience (ranging from weeks to multiple years of experience) were tasked to segment the glottal area in a random subset of the BAGLS [7] dataset (Overview Fig 2a). Experts manually annotated each frame three times in a randomized order to also gain an intra-rater reliability measure. For evaluation, we use the Intersection over Union (IoU), a common metric to compare two segmentation masks on a pixel-wise level (see Methods and [22]).

Figure 2 shows an overview of the intra- and inter-rater reliability (IaRR and IeRR, respectively). We found that the average IaRR is on average 0.77 *±* 0.01 (mean *±* std), whereas the IeRR is 0.70 *±* 0.06 as derived from Fig 2b and Fig 2d, respectively. This is in line with previous studies investigating the inter-segmenter reliability [7]. We additionally investigated how IaRR and IeRR change relative due to the average amount of labeled pixels and the distance to the center of mass. As shown in Fig. 2f, we found that especially small areas (lower than 20 px) have a low IaRR and IeRR (0.029 *±* 0.042 and 0.26 *±* 0.00), respectively) compared to rather large areas (more than 200 px), showing very high IaRR and IeRR values (0.81 *±* 0.01 and 0.66 *±* 0.07, respectively). IeRR and IaRR are also highly consistent close to the center of mass, but decrease dramatically when plotted relative to the edge. In addition, Figure 2e shows a histogram illustrating the intra-rater reliability (IaRR) across six raters. This visualization highlights a trend where the frequency of IoU scores tends to increase with higher IoU values, indicating a tendency towards greater annotator consistency at higher levels of agreement. Furthermore, we also investigated the inter-model reliability (similar as IeRR) for previously established glottis segmentation networks [8]. These networks are contained in the GAT software and are established segmentation tools, namely a custom-tailored U-Net (*Fast*, similar to [9]) and two U-Nets based on the ResNet50 (*Quality*, [17]) and EfficientNetB0 (Balanced, [18]). These networks were also trained on the full BAGLS dataset (see Methods) and evaluated the same 100 frames as the raters. We found that on average the agreement is 0.788, larger than the human IaRR and IeRR, indicating higher self-consistency. Nevertheless, these value ranges are similar to the human corresponding part indicating that glottis segmentation has some degree of uncertainty for humans and machines.

### Comparison of Segmentation Quality and Intra- and Inter-Rater Reliability

Our primary objective is to compare the results of human raters and neural network models to determine which is superior in terms of IoU. To address this, we aimed to regress the IoU score using a deep neural network. First, we asked which information is crucial to accurately predict the IoU score. We systematically changed the input to the neural network as shown in Figure 3a: We either used only endoscopic images as input (eee), the segmentation mask (sss), or a combination of both (see and sse). Among these, the “see” configuration utilizing MobileNetV2 stands out as a good choice, as observed in 3b, c. Finally, we compare the RMSE of different GAT neural networks and the average of six human raters. As illustrated in 3d, EfficientNetB0 outperforms the other models, showing an overall lower RMSE indicating that its predicted IoU values are closer to the actual IoU values. In most cases, EfficientNetB0 demonstrates higher IoU values, especially in the see combination, where near-perfect quality results in higher expected IoU. Conversely, for combinations with lower quality, where a high IoU is not expected, EfficientNetB0’s predictions appropriately reflect this.

### Traffic Light System for IoU Prediction

Focusing on the EfficientNetB0 model, we observe that it predominantly features green throughout the videos 4b, consistently predicting high IoU scores where high-quality segmentation matches the ground truth. This observation is consistent with our findings where EfficientNetB0 exhibited the lowest RMSE among all models tested, in 3d, confirming its superior accuracy in predicting segmentation quality. Comparatively, other models display a significant amount of yellow and red. This pattern suggests their comparatively lower performance, especially in frames challenged by adverse conditions such as motion blur or inadequate lighting. This direct visual comparison underscores EfficientNetB0’s effectiveness in practical scenarios, particularly in accurately handling frames suitable for endoscopic analysis.

## Discussion

In this study, we showed the significance of our findings in the context of glottal area segmentation, focusing on both inter-rater and intra-rater reliability, the prediction of segmentation quality, and finally providing the traffic light system for visualizing IoU scores and assigning each frame to a specific group. Our study focused on the evaluation of glottal area segmentation, a critical aspect for the quantification of vocal fold physiology using laryngeal high-speed videoendoscopy. The predicted segmentation quality, as represented by the IoU metric, demonstrated parity with human raters. This alignment underscores the potential of our fully automatic system to reliably assess glottal area segmentation, which is important for a labor-free clinical implementation of high-speed footage quantification. The evaluation of inter- and intra-rater reliability shed light on manual annotations’ consistency and variability. Our findings are extending and confirming prior work on inter-rater reliability. An early assessment has been performed in [7], where up to four segmenters and their inter-rater reliability have been assessed. In exploring intersegmenter variability in Glottal Analysis Tools (GAT) measures, [13] conducted a cohort study assessing rater reliability. This study focused on the analysis of the glottal area waveform (GAW) from high-speed videoendoscopy, involving trained segmenters who independently annotated videos. The study’s use of the Intraclass Correlation Coefficient (ICC) adds a quantitative dimension to the evaluation of GAW measures. With high ICC values across various parameters, the study highlights the clinical applicability of GAT, indicating strong inter-rater reliability.

In line with these findings, we observed a stronger IaRR compared to the IeRR indicating that there is a self-contained ability to perform manual glottis segmentation. Nevertheless, these IeRR values are far from optimal. Examining IeRR across pixel sizes and distances to the center of mass of the glottis segmentation revealed further insights. Smaller areas (less than 20 pixels) presented challenges, exhibiting lower inter- and intra-rater reliability, most likely due to the relative high fraction of fuzzy and dubious contour pixels in small segmented areas. As the variance is 0.1 for 75 percent of the data when using EfficientNet (as seen in Figure 3), predictions that are close to the traffic light thresholds are particularly prone to misclassification. For example, if the true IoU value is 0.65, the model predicts in the range of 0.55 to 0.75 due to the observed variance. Thus, there is a high probability that predictions in this critical range will be misclassified. Misclassifications in this range can lead to moderate segmentations being considered good and vice versa. This affects the reliability of the automatic segmentation assessment and can lead to incorrect medical decisions being made. This also means that very poor segmentations and very good segmentations are reliably identified by our model, resulting in a very low misclassification rate in these regions.

Our investigation into predicting segmentation quality through IoU score regression revealed dependencies. Systematically altering neural network inputs, including endoscopic images, masks, or their combination, helped us in fine-tuning the system’s IoU. The traffic light system, visually representing IoU scores, demonstrated the influence of pixel size and expert-derived scores on prediction accuracy. Our analysis included comparisons between human raters and various neural network models such as CustomUnet Fast, ResNet50 Quality, and EfficientNetB0 Balanced. While all models demonstrated competitive capabilities, the EfficientNet model slightly outperformed others, including human raters, in terms of RMSE and showed a predominance of green in the traffic light visualization, indicating higher segmentation quality consistently across frames. The successful alignment of automated predictions with human raters supports our system’s integration into clinical workflows.

The traffic light system offers a user-friendly interface for clinicians, promoting a collaborative human-in-the-loop approach. This model, merging AI strengths with expert judgment, holds promise for enhancing glottal area segmentation reliability and efficiency in clinical settings. Overall, this study underscores the benefits of AI in enhancing diagnostic procedures in laryngeal endoscopy and sets the stage for further innovations in medical imaging. Our traffic light system for visualizing IoU scores provides an easy-to-grasp, quick evaluation framework for downstream analysis. However, we observed that especially small segmentation areas have a high variance in IoU prediction. Future work should look into refining the algorithm to better handle these small segmented areas, which currently show lower reliability, consistent with previous reports [7]. The use of more elaborate architectures may yield better outcomes as suggested in recent studies [23, 24]. Improving the system’s ability to accurately analyze these small segments could significantly enhance overall accuracy.

## Conclusion

In conclusion, our study dives into the details of glottal area segmentation, reliability among raters, and predicting the quality of segmentation in laryngeal endoscopy. Our findings also shed light on the inter-rater reliability among human annotators and neural network models. Notably, while neural networks generally perform well, they are not without issues. One possible reason for discrepancies in neural network performance could be the inherent variability in human annotations used for ground truth data. This variability might affect the learning outcomes of the models, potentially leading to inconsistencies in segmentation quality predictions.

This performance edge of the EfficientNet underscores its potential to supplement human judgment in clinical settings. The close results among different models, such as CustomUnet, ResNet50, and EfficientNetB0, compared to human raters, indicate the critical role of high-quality, consistent annotations in training robust models. As we continue to refine our techniques and expand our model comparisons, our goal is to enhance the reliability of both human and automated assessments. This effort will not only improve the accuracy of diagnostics but will also support the broader integration of AI tools in healthcare and effective patient care through advanced technology.

## Supporting information

N/A

## Acknowledgments

We thank Marion Dörrich, Patrick Schlegel, Hernan Aguilera for technical support and data annotations.

## Footnotes

1 https://github.com/pvigier/perlin-numpy

